# ICP1 bacteriophage treatment antagonizes colonization of the zebrafish larval intestine by *Vibrio cholerae*

**DOI:** 10.1101/2025.09.01.673471

**Authors:** Adam Sidi Mabrouk, Jamie S. Depelteau, Chiara Foini, Annabel Kempff, Sebastiaan Jonker, Susanne Brenzinger, Ronald Limpens, Manuel Majrouh, Annemarie H. Meijer, Ariane Briegel

## Abstract

Outbreaks of cholera pose a major threat to human health. Currently, antibiotics are the most effective treatment against the causative agent, the bacterium *Vibrio cholerae*. However, the use of antibiotics eventually leads to the emergence of resistant strains, which necessitates the need for alternative approaches. The use of bacteriophages to target the infection by antibiotic resistant bacteria is one promising alternative. While clearance of *Vibrio cholerae* with the use of phages has been performed on several animal models, none of these models are natural hosts of *V. cholerae*. Therefore, we set out to investigate the interaction between *V. cholerae* and bacteriophage ICP1 both *in vitro* and *in vivo* in a natural host, the zebrafish model, *Danio rerio*. To study the interplay between host, bacteria and phages we used a combination of light and ultrastructural imaging techniques, including confocal fluorescence microscopy, serial block face scanning electron microscopy (EM) imaging and cryogenic EM, which allowed us to investigate both the colonization process by *V. cholerae* and clearance by the ICP1 bacteriophage. In addition, we determined the effects of the microbiome on this treatment by using germ-free, conventionalized and monoassociated zebrafish larvae as a host. Independent of the presence and composition of microbiomes used here, *V. cholerae* efficiently colonized the larval intestine. Finally, we demonstrate significant in vivo clearance of V. cholerae N16961-dsRED by ICP1, underscoring the role of phage-bacteria dynamics in shaping pathogen colonization within the zebrafish larval host.

**SIGNIFICANCE:** Cholera remains a life-threatening disease that causes recurring outbreaks and significant mortality, particularly in developing and conflict-affected regions. As antimicrobial resistance continues to rise, there is an urgent need to better understand the ecological and microbial dynamics that govern *Vibrio cholerae* colonization and persistence. This research investigates how *V. cholerae* interacts with bacteriophages, the host environment, and the resident microbiota within a natural vertebrate host, offering new insights into the factors that influence pathogen clearance and shaping of the gut ecosystem during infection. The powerful combination of serial block-face scanning and cryogenic electron microscopy, fluorescence microscopy, and traditional colony/plaque counting methods revealed previously unobserved aspects of the interplay between host, pathogen, phages, and the microbiome, highlighting phage-driven clearance of *V. cholerae* during colonization.

## INTRODUCTION

The water-borne bacterium *Vibrio cholerae* is the causative agent of the diarrheal disease cholera. It remains a major threat to human health with the most recent pandemic causing up to 2.86 million cases and an estimated 95.000 of these cases resulting in death (1). Unlike the other six pandemics before it, the seventh pandemic is caused by *V. cholerae* of the El Tor biotype (2). Clinical interventions that actively target the bacteria are mostly limited to the use of antibiotics. While the use of antibiotics shows a decrease in the burden of cholera (3), antibiotic resistant strains have already emerged (4). Therefore, there is an increasing interest in developing alternative treatments, such as the use of bacteriophages.

Bacteriophages, viruses that specifically infect bacteria, offer a promising therapeutic strategy due to their host specificity and self-limiting life cycles (5, 6). Lytic phages, in particular, rapidly replicate within and lyse bacterial cells without integrating into the host genome, minimizing the risk of horizontal gene transfer of virulence or resistance determinants (7). This makes lytic phages appealing candidates for antimicrobial treatment.

Interest in bacteriophages as a therapeutic tool against bacterial infections dates back over five decades (7), and recent research has reignited this interest, particularly in the context of combating antibiotic-resistant pathogens (5, 8, 9). Phage therapy studies have relied heavily on mammalian models, such as rabbits and infant mice, to evaluate the efficacy of phages against *Vibrio cholerae* infections. In these models, oral administration of *V. cholerae*-specific phages has been shown to reduce disease severity—marked by decreased diarrhea—and lower bacterial loads in feces, as measured by colony forming units (CFUs) (10). Similar results were observed with a prophylactic administration of a phage cocktail containing phages ICP1, ICP2, and ICP3 (11). Here, they also observed a reduction in the amount of enumerated *V. cholerae*. Although resistant bacterial variants emerged within hours, these phage-resistant strains exhibited reduced pathogenicity, suggesting a fitness trade-off.

While previous studies have demonstrated the effectiveness of bacteriophage therapy in various animal models, these systems often rely on mammalian models which are not exposed *V. cholerae* in natural environments. This is reflected in the surgical interventions which are necessary to allow *V. cholerea* to colonize the intestine in the RITARD (removable intestinal tie-adult rabbit diarrhea) rabbit model (12). In contrast, zebrafish (*Danio rerio*) larvae naturally ingest *V. cholerae* from their surrounding aquatic environment, allowing colonization to occur without experimental manipulation (12). Furthermore, zebrafish larvae represent a more ecologically relevant host, sharing freshwater habitats with environmental *V. cholerae* populations in South Asia, where the bacterium is endemic (12, 18, 19). Moreover, zebrafish larvae are translucent, making them especially suited for the use of light microscopy. This positions zebrafish larvae as a practical infection model to study the interactions between *V. cholerae* and bacteriophages with which it shares an aquatic niche. By studying these interactions in a natural host, we can better understand how *V. cholerae* persists in a natural environment.

In this study, we show the effects of ICP1 bacteriophage treatment on *V. cholerae* both *in vitro* and *in vivo*. The *in vitro* results indicate that the efficacy of bacteriophage predation of *V. cholerae* by ICP1 bacteriophage predation is dependent on the aqueous environment. Furthermore, by imaging the colonization of the intestine of zebrafish larvae by *V. cholerae*, of the El tor biotype, with confocal fluorescence microscopy, cryo-EM and serial block face scanning electron microscopy (SBF-SEM), we gained insight on the colonization dynamics of the bacteria in the presence and absence of microbial communities. Following administration of ICP1 to the *V. cholerae* infected larvae, a significant decrease in bacteria in the intestine was observed and quantified by selective plating. Combined, these results elevate the zebrafish larvae as a natural host for pandemic *V. cholerae* and demonstrate the effectiveness of the ICP1 bacteriophage in clearing colonization within the host.

## RESULTS

### Interaction between *V. cholerae* and ICP1 *in vitro* depends on the aquatic environment

In order to investigate the interactions between *V. cholerae*, the zebrafish larvae and the bacteriophage, we selected the widely used bacteriophage ICP1, as it has been naturally found in Bangladesh waters, where zebrafish also naturally occur (22, 23). In addition, it has previously been isolated from patient stool samples, highlighting its clinical relevancy (24). To minimize the complexity of experimental conditions while investigating the interaction between bacteria and phage, we first studied their interaction in the absence of the zebrafish host. This allowed us to gain insight into the influence of the aquatic environment itself on the phage infection. More specifically, we aimed to understand if different types of lab media would affect the interaction compared to a natural water source (Rhine water). Therefore, we investigated *V. cholerae* N16961-DsRed predation by ICP1 in multiple environmental conditions outside of the host. First, we used standard rearing media of zebrafish larvae, artificial fresh water (AFW) as this will be the aqueous environment in which the larval infection experiments will take place. As a control, we repeated the experiments in LB media, which is the standard growth media for *V. cholerae* (25). As expected, we observed the clearance of *V. cholerae* after 2 hours in LB media, which also coincided with an increase in the number of bacteriophages up to a PFU of 10^9^ bacteriophages per mL (Fig. 1A). The clearance continued until the 7-hour timepoint, afterwards we observed the resistance of *V. cholerae* to ICP1 as can be seen by an increase in CFU. However, when repeating these conditions in AFW, we observed no clearance of *V. cholerae* by ICP1 and no propagation of bacteriophages was detected (Fig. 1A). These results imply that phage predation is limited in the AFW environment.

**Figure 1.**
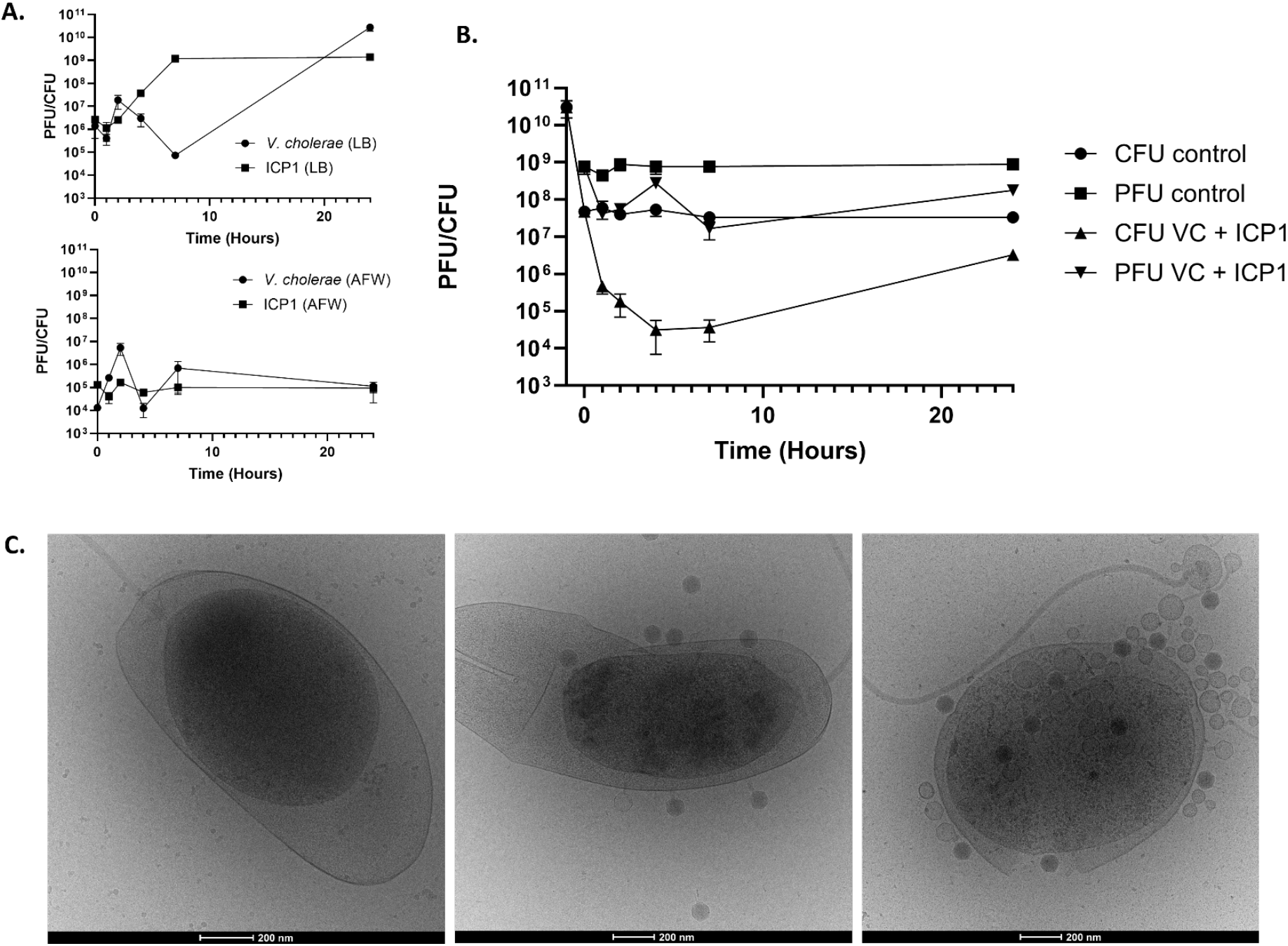
Interaction of *V. cholerae* and ICP1 in various saline environments. (A) Graphs depicting the interaction between *V. cholerae* (CFU; circles) and ICP1 (PFU; squares) in LB and AFW over time. The phage was added to these cultures at timepoint 0 at a multiplicity of infection (MOI) of 2. These are the results of three individual experiments for each treatment. Error bars represent standard error of mean (SEM). (B) Graph depicting the interaction between *V. cholerae* and ICP1 in natural Rhine water after pre-incubation in the water. The -1 hour timepoint represents concentration of bacteria before preadaptation The phage was added to these cultures at timepoint 0 hours at a multiplicity of infection (MOI) of 2. These are the results of three individual experiments for each treatment. Error bars represent standard error of mean (SEM). (C) Cryo-electron micrographs of preconditioned *V. cholerae* with ICP1. Representative images from timepoints 0 (left) and 24 hours (middle and right) are depicted.

While the AFW is a standardized environment that is widely used to rear zebrafish in laboratory conditions, it does not completely mimic a natural water source. Therefore, we investigated predation by ICP1 in natural freshwater conditions using water samples collected from the Rhine river. To mimic the natural growth conditions as closely as possible to a previously published study (26), *V. cholerae* was also preadapted by growing the culture overnight in freshwater. When repeating the experiments in this media, we could clearly observe a drop in the number of bacteria, coinciding with an increase of bacteriophages (Fig. 1B). This indicates that ICP1 can propagate and lyse *V. cholerae* under these conditions. Closer inspection of these samples through cryo-electron microscopy revealed that *V. cholerae* displayed a detachment of the inner membrane and shrinkage of the cytoplasm, also referred to as plasmolysis, of the inner membrane at all time points (Fig. 1C). Curiously, this still led to the binding and even propagation of bacteriophage ICP1. While the observed plasmolysis is likely the cause of the decrease in bacteriophage propagation, it does not completely abolish bacteriophage infection in natural freshwater. In conclusion, our data shows that natural freshwater allows the predation of *V. cholerae* by ICP1 while the standardized zebrafish rearing media, AFW, does not.

### *V. cholerae* shows robust colonization of germ-free zebrafish larvae

Having established a baseline understanding of how *V. cholerae* interacts with the ICP1 bacteriophage in different types of media, we wanted to gain insight into how the interaction between phage and bacteria unfolds inside the zebrafish host. The fish were reared in AFW as this is a controlled environment that is widely used in research laboratories (27). First, we needed to confirm that our bacterial laboratory strain *V. cholerae* N16961-DsRed robustly colonizes the fish intestine. To do this, we performed infection experiments in germ-free zebrafish larvae. The absence of a natural gut microbiome reduces the complexity and allows for clear visualization of the colonization of *V. cholerae* N16961-DsRed. At 3 days post-fertilization (dpf) one group of germ-free larvae was infected with 10^9^ bacterial cells of *V. cholerae* N16961-DsRed. This was done by adding the inoculum directly to the water of the zebrafish larvae. The bacteria are then passively ingested by the larvae, eliminating the need for injection. The other group was uninfected and acted as the control group. We investigated the colonization of 5 dpf using confocal fluorescence microscopy, 48 hours after the addition of *V. cholerae*. Z-stack series were generated at two distinct locations in the intestine (Figure 2A; S1; Movie S1). One area is located directly below the swim bladder with the other area moving towards the distal end of the anterior intestine and into the mid-intestine. Red fluorescent bacteria were observed in both areas of the intestine, indicated by a higher fluorescence intensity compared to the background fluorescence of the intestine in the control larvae (Figure 2A & S1.) Furthermore, both microcolonies and bacteria that were free flowing through the intestine were observed (Figure 2A; Movie S2). This confirmed that a significant amount of *V. cholerae* N16961-DsRed cells is able to colonize the anterior intestine of the germ-free zebrafish larvae.

**Figure 2.**
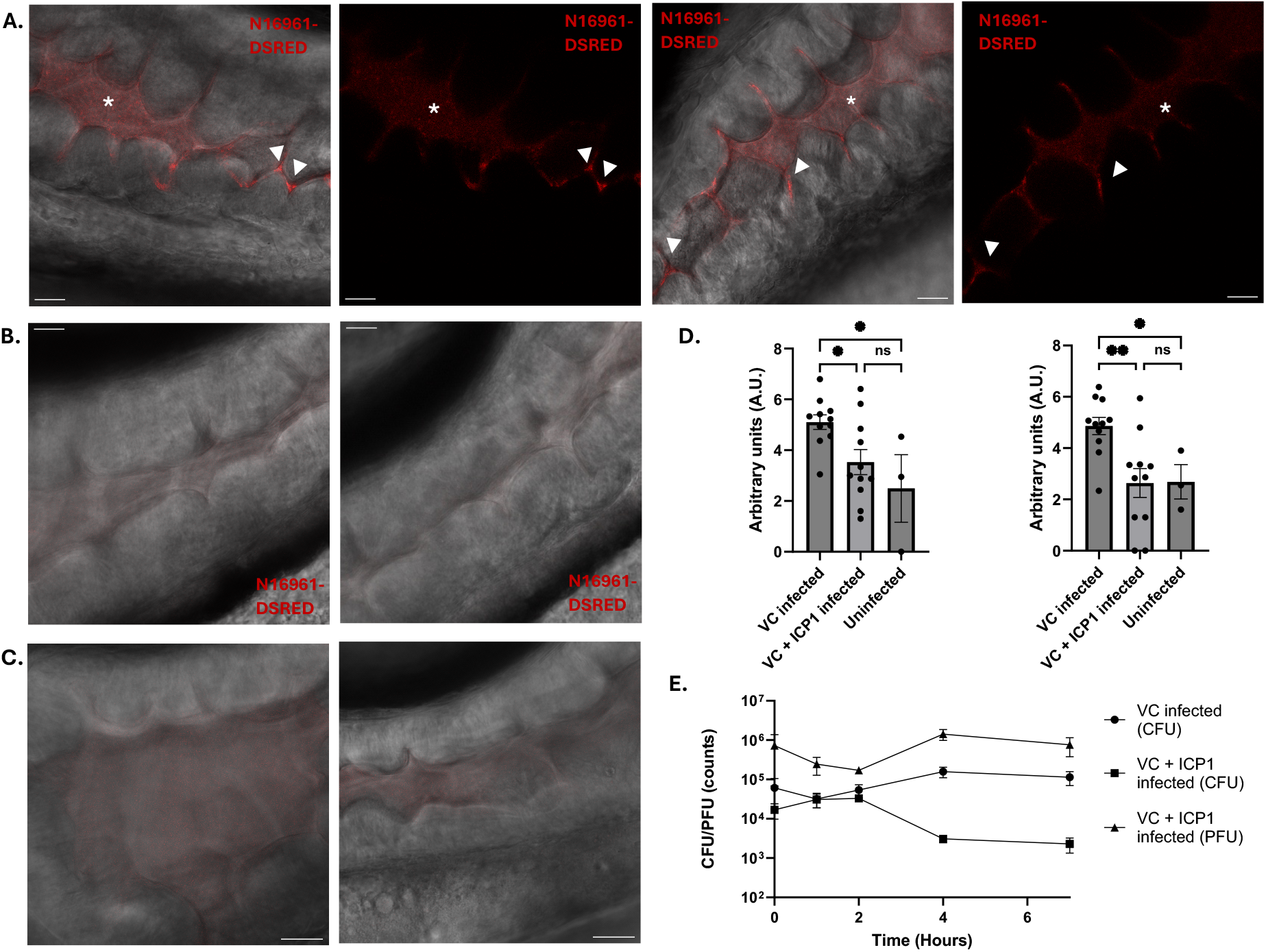
Clearance of *V. cholerea* N16961-DSRED infection by ICP1 in germ-free zebrafish larvae. Single slice images of confocal fluorescence Z-stacks showing the anterior (left) and mid-intestine (right) of germ-free zebrafish larvae either infected by *V. cholerae* (A); by *V. cholerae* and ICP1 (B) or uninfected (C). Composite images of bright field and fluorescent signal are shown. Clumps of microcolonies (white arrowheads) are visible in the crevices between the villi of the intestine at both locations. Small specks of red fluorescent free-floating bacteria are present throughout the anterior intestine and are most commonly found in the open areas of the intestine (white asterisk). Scale bars represent 20 μm. (D) Graphs depicting the log-normalized fluorescent pixels that were quantified of colonizing bacteria in the anterior (left) and mid-intestine (right). Error bars represent standard error of mean (SEM). An ordinary one-away ANOVA was performed for the comparison of means (^*^P < 0.05, ^**^P < 0.005). These are the results of three independent experiments with 11 larvae for the *V. cholerae* and *V. cholerae* + ICP1 infected groups and 3 larvae for the uninfected group. (E) Graph depicting the amount of CFU and PFU counts per larvae in untreated (circle) and treated larvae (square; triangle) over time. Error bars represent standard error of mean (SEM). Each individual point is the average of each replicate; with each replicate being the average of 5 fish. Three independent experiments were performed in total.

To investigate the colonization of the zebrafish larvae in greater detail, we employed serial block face imaging, a technique that allows the inspection of the morphology and localization of the bacteria inside of the zebrafish intestine. Here, we compared germ-free 5 dpf zebrafish larvae both infected and uninfected by *V. cholerae* C6706-tdTomato. Using this method, we were able to achieve large volumes of three-dimensional data covering up to 150 μm length of the intestine. An overview of the uninfected zebrafish larvae allowed us to confirm the lack of other microbes in the anterior intestine and observe free floating debris and the loose packing of the villi (Fig. 3A). In contrast, the microvilli of the *V. cholerae*-infected larvae were tightly packed, and microcolonies were present throughout the lumen, including the base of villi as well as unassociated with the intestinal lumen (Fig. 3B & Fig. 3C, arrowheads). Imaging of the larvae by confocal microscopy also showed that the infection of the intestine is similar to that observed with the N16961-dsRED strain (Fig. S2). We also observed that the infection decreases as it approaches the mid- and posterior intestine, areas where the majority of the mucus producing goblet cells reside (Fig. 3C; (28)). Furthermore, the three-dimensional volume images allowed the analysis of the overall morphology of the bacterial cells that colonized the intestine. Using segmentation software (Dragonfly; (29)), we were able to outline part of the anterior intestine and the bacteria within (Fig. 3D; Movie S3). Here we can see the abundance of the bacteria in the larval intestine. This also allowed us to clearly identify colonizing bacteria in the base of the villi, including some dividing bacteria (Fig. 3E). Overall, all cells exhibited a vibrioid shape (Fig. 3), which contrasts with the more spherical morphology in cells exposed to the Rhine water (Fig. 1C). Together, these observations confirm that pandemic *V. cholerae* is capable of robust colonization of the zebrafish larval intestine, with the morphology of the bacterial cells indicating that they are viable.

**Figure 3.**
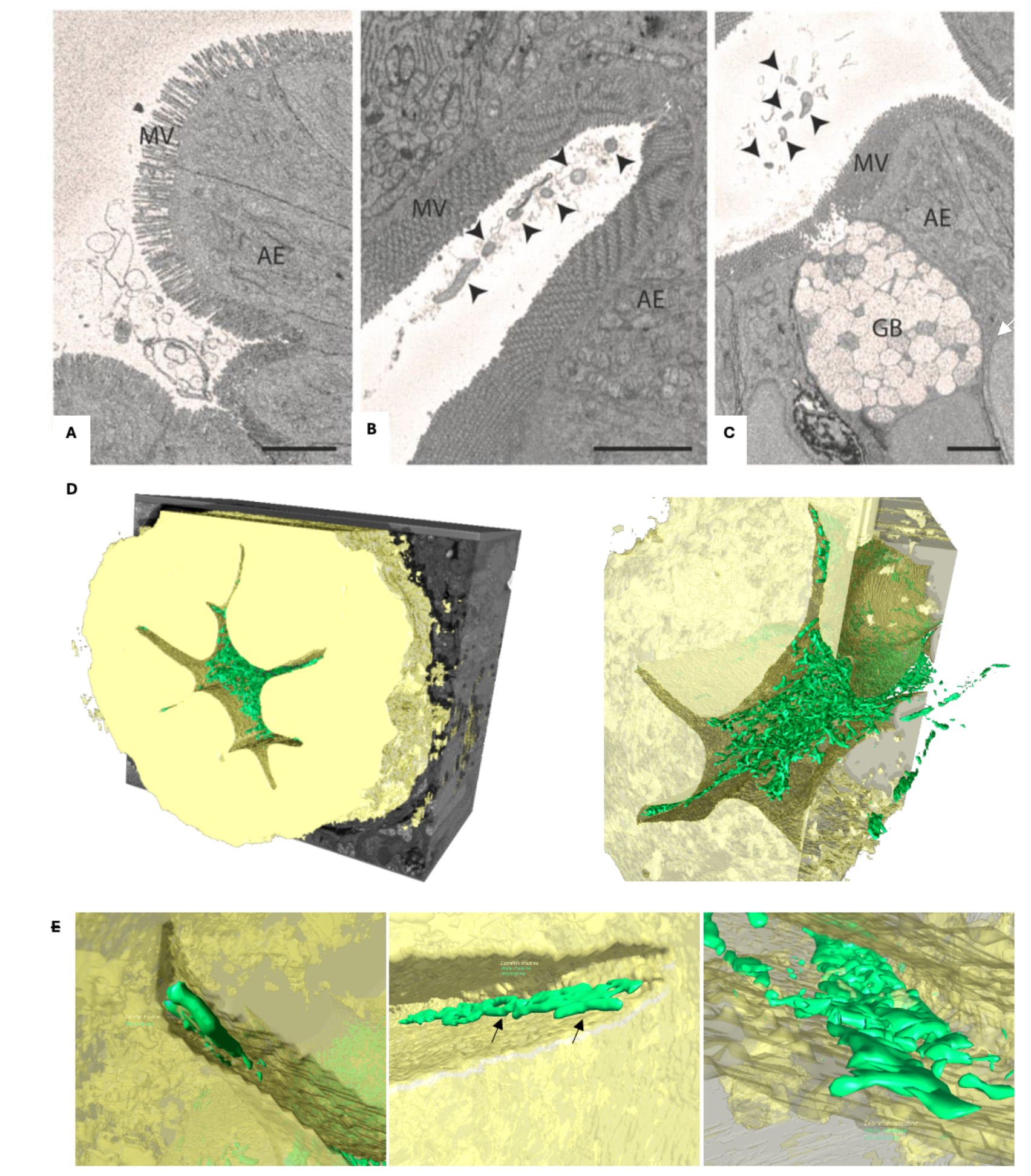
Serial block face images and segmentation of zebrafish larvae intestine both infected and uninfected by *V. cholerae* C6706-tdTomato. Representative SEM block face images of the anterior intestine of the germ-free (A) and infected (B, C) zebrafish larvae. *V. cholerae* cells are indicated with black arrowheads. (D) Overview images of the segmentation of the anterior intestine. Intestinal tissue is depicted in yellow and *V. cholerae is* depicted in green. (E) Representative segmented SBF SEM sections depicting areas of the intestinal villi which are colonized by *V. cholerae* microcolonies. Dividing bacteria have been highlighted (black arrows). SB, swim bladder; AE, absorptive enterocytes; MV, microvilli; GB, goblet cells. Scale bar = 3 μm (A-C).

### Phage-mediated clearance of *V. cholerae* occurs in both germ-free and conventionalized zebrafish larvae

Now that the colonization of the zebrafish larvae intestine by *V. cholerae* N16961-dsRED had been established, we assessed the ability of ICP1 to prey on *V. cholerae* within the host. This was done by repeating the infection experiments as previously described, but now with the addition of ICP1. The ICP1 treatment group showed very little fluorescence in both areas of the intestine and looked similar to the uninfected control intestine (Fig. 4BC; Movie S4). To visually quantify the effect of ICP1 predation, the fluorescent signal was quantified and compared between all conditions (Fig. 3D). Here, we observed that the ICP1 treated larvae have a reduction in fluorescent signal of microcolonies, both in the anterior intestine and in the mid-intestine. CFU and PFU counts of all treatments were also performed to quantify the total amount of *V. cholerae* and ICP1 present in the fish (Fig. 3E). We observed stabilization of *V. cholerae* infection after four hours, with the ICP1 treated larvae showing a clear reduction in bacterial load by a factor of 10. Which was further confirmed by imaging of larvae 4 hours after treatment with ICP1 (Fig. S3). This coincides with an increase in the number of phages, indicating that the decrease in bacterial load is caused by bacteriophage predation.

**Figure 4.**
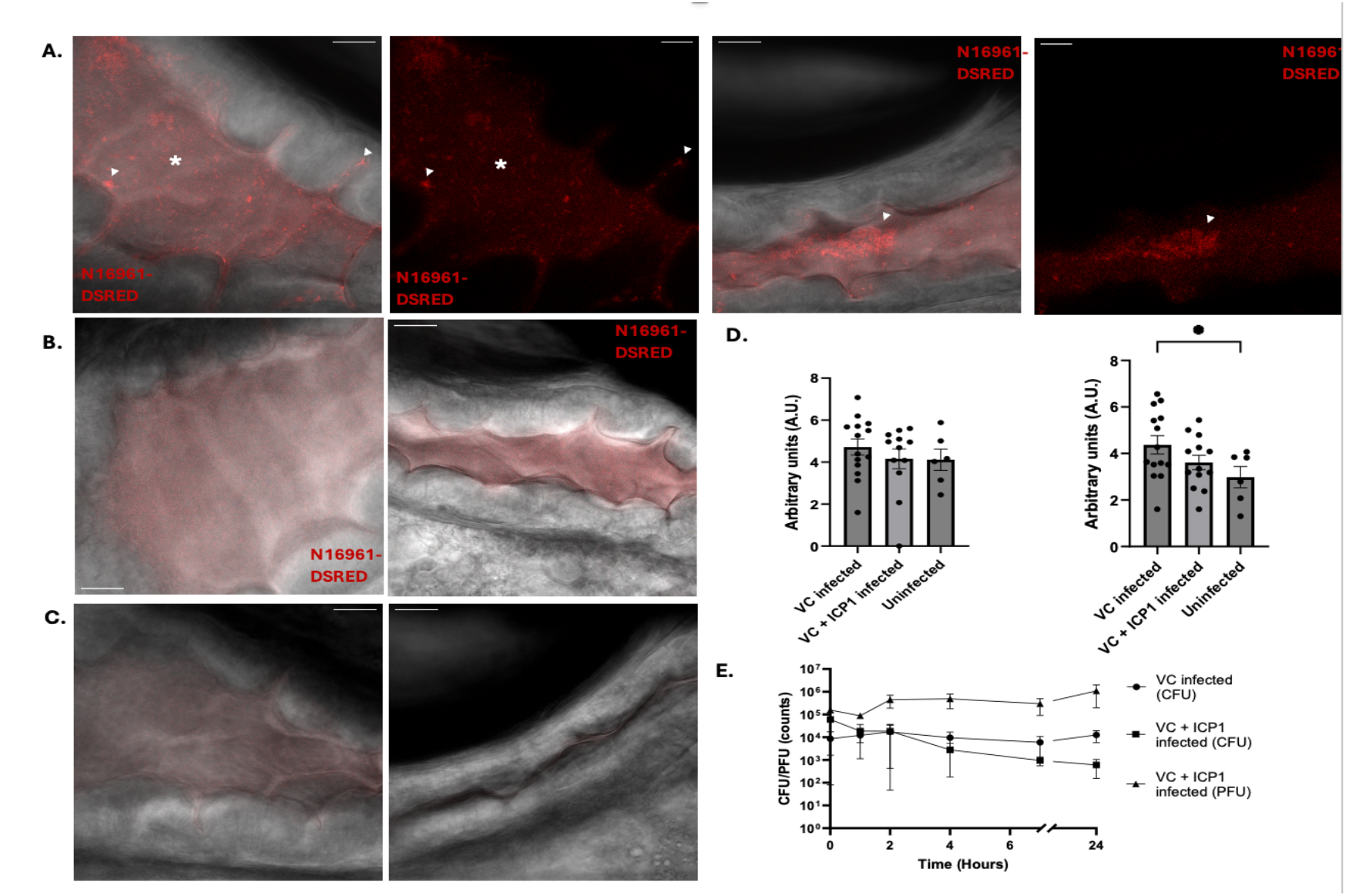
Infection and clearance of *V. cholerae* N16961-DSRED by ICP1 in conventionalized zebrafish larvae. Single slice images of confocal fluorescence Z-stacks showing the anterior (left) and mid-intestine (right) of germ-free zebrafish larvae either infected by *V. cholerae* (A); by *V. cholerae* and ICP1 (B) or uninfected (C). Composite images of bright field and fluorescent signal are shown. Clumps of microcolonies (white arrowheads) are visible in the crevices between the villi of the intestine and free floating within the intestine. Small specks of red fluorescent free-floating bacteria are present throughout the anterior intestine and are most commonly found in the open areas of the intestine (white asterisk). Scale bars represent 20 μm. (D) Graphs depicting the log-normalized fluorescent pixels that were quantified of colonizing bacteria in the anterior (left) and mid-intestine (right). Error bars represent standard error of mean (SEM). An ordinary one-away ANOVA was performed for the comparison of means (^*^P < 0.05). These are the results of four independent experiments with 14 larvae for the *V. cholerae* infected group; 12 larvae for the *V. cholerae* and ICP1 infected group and 6 larvae for the uninfected group. (E) Graph depicting the amount of CFU and PFU counts per larvae in untreated (circle) and treated larvae (square; triangle) over time. Error bars represent standard error of mean (SEM). Each individual point is the average of each replicate; with each replicate being the average of 5 fish. Three independent experimdents were performed in total.

While there is a significant amount of clearance of *V. cholerae* in the germ-free system, we also wanted to assess the ability of ICP1 to lyse *V. cholerae* in conventionalized zebrafish larvae. The microbiome could have a strong effect on both the colonization of the intestine by *V. cholerae* and the efficacy of the ICP1 treatment (30–32). Therefore, the experiments were repeated in fish with a native microbiome. When imaging the untreated larvae, we observed similar amount of colonization in both anterior and mid-intestine (Fig. 4A). Specifically, we observed large microcolonies of bacteria both free floating and in the base of the intestinal villi. Quantification of the fluorescent signal showed that the level of colonization is equally high in both parts of the intestine and is within range of the germ-free larvae (Fig. 4D & Fig. 3D). Imaging of the larvae that were treated by ICP1 showed that both the anterior area of the intestine and the mid-intestine have a lower fluorescent readout as most of the large clumps of fluorescent signal seems to be diminished (Fig. 4B). Quantification of the fluorescent signal in this setup does point towards an overall higher fluorescent readout in all conditions (Fig. 4D). The CFU and PFU counts display a drop of almost 10-fold in the number of bacteria, following an increase in bacteriophages between 1 – 4 hours after treatment by ICP1 (Fig. 4E). Notably, this drop remained consistent after 24 hours of ICP1 treatment (Fig. 4E). Taken together, the data indicates that the clearance of *V. cholerae* by ICP1 is slightly reduced in conventionalized larvae but consistent over a longer period of time.

### Monoassociation of *Aeromonas sp*. or *Pseudomonas sp*. does not affect colonization and clearance of *V. cholerae*

While the ICP1 mediated clearance remained effective in conventionalized zebrafish larvae, the CFU and PFU counts showed higher variability than in the germ-free system (Fig. 4E), pointing towards an influence of the microbiota on the colonization and predation by ICP1. Therefore, we set out to identify the main members of the microbiota in the zebrafish larvae intestine within this setup. For this purpose, we plated the water of the larvae every day during the infection process on TCBS plates, which revealed two main colony morphologies (Figure S4). By performing 16S RNA sequencing, we identified that these colonies are *Aeromonas sp. (yellow)* and *Pseudomonas sp. (green)*. While these plates are normally only selective for *Vibrio* species, the detection of other bacterial species is in line with a previous report (33). Next, we repeated the previous infection experiments with germ-free larvae, but this time they were now monoassociated with either *Aeromonas sp*. or *Pseudomonas sp*. for 30 minutes before the addition of *V. cholerae*. Qualitative imaging of monoassociated larvae showed similar results to our previous findings. Namely, we observed robust colonization of the intestine with subsequent clearance after addition of ICP1 (Fig. 5AB). This is also reflected in the CFU/PFU counts, where we observed both an increase in the number of phages after 2 hours and a decrease in bacteria at the same time, while the number of bacteria remained stable in the untreated larvae (Fig. 5D). This indicates that the presence of either of these bacterial species does not affect the colonization or subsequent treatment of the colonization by the ICP1 bacteriophage.

**Figure 5.**
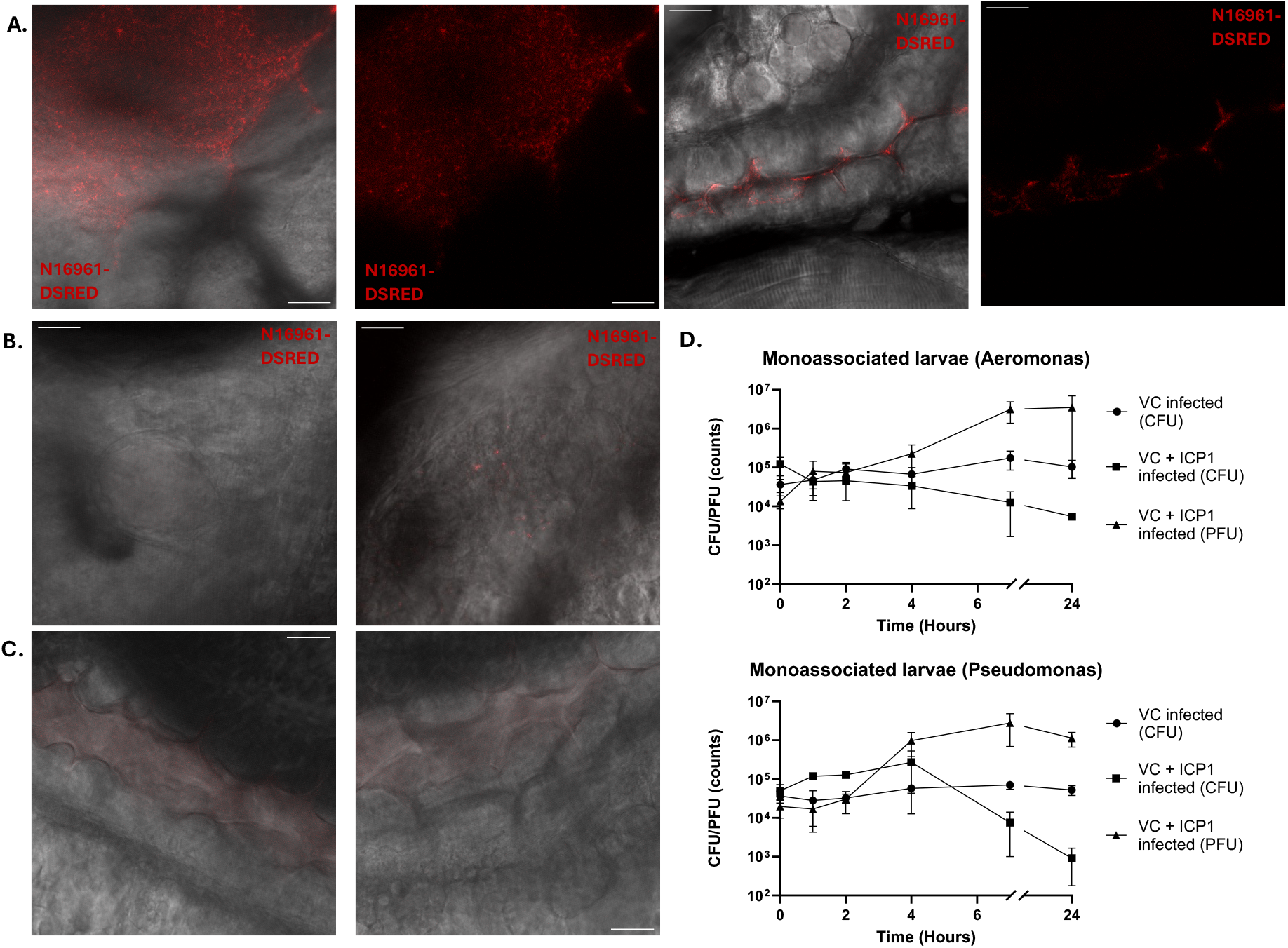
Infection and clearance of *V. cholerae* N16961-DSRED by ICP1 in zebrafish larvae monoassociated by either *Aeromonas sp*. or *Pseudomonas sp*. Single slice images of confocal fluorescence Z-stacks showing the anterior intestine of zebrafish larvae monoassociated by *Aeromonas sp*., infected either by *V. cholerae* (A); *V. cholerae* and ICP1 (B) or uninfected (C). The anterior intestine was imaged directly below the swim bladder (left) and at the mid-intestine (right). Scale bars represent 20 μm. (D) Graphs depicting the CFU and PFU counts in untreated (circle) and treated larvae (square; triangle) over time. Error bars represent standard error of mean (SEM). Each individual point is the average of each replicate; with each replicate being the average of 5 fish. Three independent experiments were performed in total.

## DISCUSSION

Understanding how bacteriophages influence colonization dynamics within a natural host is critical to unravelling the interactions that shape the persistence and spread of *Vibrio cholerae*. Here we show the clearance of *V. cholerae* colonization by ICP1 bacteriophages in a zebrafish larvae model with either a devoid, intact, or monoassociated intestinal microbiota. The combination of confocal fluorescence microscopy, SBF SEM, and colony counting yields new insights into the colonization dynamics over time and subsequent clearance by ICP1. These findings reveal how *V. cholerae* colonization and clearance are influenced by the presence or absence of the intestinal microbiota and demonstrate the capacity of ICP1 phages to impact pathogen dynamics within a vertebrate host. Furthermore, by showcasing the ability of pandemic *V. cholerae* strains to effectively colonize the zebrafish larvae intestine we provide a basis for an ecological niche that the zebrafish larval intestine forms for *V. cholerae* and the propagation of bacteriophages.

By examining the lifecycle of *V. cholerae* and ICP1 in different media, we observed how environmental conditions influence bacterial growth and phage propagation. In artificial freshwater (AFW), *V. cholerae* exhibited minimal growth even without phages, unlike in LB. Interestingly, this contrasts with our in vivo zebrafish infections, where bacterial loads increased approximately tenfold, likely due to higher nutrient availability in the intestine (34, 35). The poor growth in AFW may explain reduced phage propagation, as nutrient limitation can impair phage propagation. This difference may be due to higher ion concentrations (e.g., Ca^2+^, Mg^2+^) in Rhine water (36), which enhance phage activity, or due to pre-adaptation of *V. cholerae* to this environment prior to infection (26). Notably, while pre-adaptation did not fully prevent plasmolysis, an indicator of nutrient stress (37), cryo-EM and plaque assays confirmed bacteriophage binding and phage propagation. Nevertheless, predation of *V. cholerae* remained largely unaffected by plasmolysis, as we observed lysis of *V. cholerae* and an increase in bacteriophages. These findings highlight the importance of environmental context in shaping host–phage dynamics.

Serial block-face imaging provided valuable insights into *V. cholerae* colonization of the zebrafish larval gut under sterile conditions. We observed bacterial presence throughout the intestinal lumen, with cells exhibiting the characteristic comma-shaped morphology—indicative of a metabolically active state (38)—and evidence of active division. Microcolonies were detected both free-floating and closely associated with the microvilli. Importantly, using a combination of SBF SEM and confocal imaging, we confirmed that two distinct clinical isolates from the ongoing seventh pandemic, C6706 and N16961, are both capable of robustly colonizing the zebrafish intestine. These strains, which belong to the O1 El Tor biotype (38, 39), successfully established themselves within the host, suggesting that pandemic *V. cholerae* strains can exploit the larval zebrafish intestine as a compatible colonization niche.

The germ-free zebrafish larvae model is a powerful tool for the investigation of host microbe interactions, due to the lack of other bacteria that could affect the host-microbe interaction. Indeed, we observed that this model benefits confocal fluorescent imaging, as the absence of the gut microbiome results in a clearer fluorescent signal and less background fluorescence. However, a fully intact gut microbiome has not only been shown to be imperative for the immune response of zebrafish larvae (40), but also provides direct interference with the colonization of pathogens (41). Interestingly, we found only minor effects on the colonization by *V. cholerae* compared to the germ-free larvae. While fluorescent quantification implies that ICP1 predation is heavily impaired in the conventionalized larvae, the reduction of large clumps of microcolonies in the images and the CFU/PFU counts indicate that ICP1 is still actively lysing *V. cholerae*. These colonizing microcolonies have been suggested to be critical for pathogenesis and persistence (42). Starting from 4 dpf, the zebrafish larvae exhibit a differentiated intestine with differences in cell types and morphology at anterior and mid to posterior positions (43). A prominent difference is that the anterior intestine has multiple larger folds, while the mid to posterior intestine remains mostly smooth with very little folds (44, 45). We considered that this may impact the ability of *V. cholerae* in forming robust microcolonies and may result in higher susceptibility to predation by ICP1. However, we observed that ICP1 treatment reduced colonization in both areas of the intestine, which highlights the potential of phage therapy as a treatment for infections.

Although colonization of the zebrafish larval intestine by *V. cholerae* was generally similar between germ-free and conventionalized conditions, we observed slightly reduced colonization and more variable clearance by ICP1 in the conventionalized larvae, suggesting that components of the native microbiota may influence both pathogen establishment and phage predation dynamics. To determine which bacteria were overly present in the microbiome, and could be the cause of interference, we employed a crude approach of selective plating and 16S sequencing. The presence of unculturable or slow growing bacteria will not be selected in this manner, however slow growing bacteria specifically would likely exert less of an effect on the colonization and predation of *V. cholerae*, due to their low abundance and the short timeframe of the larval infection experiment. Through this approach, we identified *Aeromonas sp*. and *Pseudomonas sp*, which is in line with microbes that have previously been found in zebrafish facilities and on TCBS selective plates (46, 47). The isolation of these strains allowed us to perform mono-associations of germ-free larvae in order to assess phage predation in a better defined microbial environment. *Aeromonas sp*. or *Pseudomonas sp*. exerted only a small effect on the colonization and clearance of *V. cholerae* in the zebrafish larvae, demonstrating that these strains only have a minor influence on the clearance of *V. cholerae* by ICP1.

Our findings raise the possibility that pandemic *V. cholerae* and its phage ICP1 may transiently occupy a shared environmental niche within aquatic vertebrate hosts such as zebrafish larvae. Because of this it is important to understand how *V. cholerae* and ICP1 interact within such a setting. Various reports state that the efficacy of phages diminish over time, as bacteria develop resistance to phage predation (48). While *in vitro* experiments revealed a resurgence of bacterial growth after 24 hours of ICP1 exposure, this pattern was not observed *in vivo*, where *V. cholerae* levels remained low or declined further. This suggests that the host environment imposes constraints on the development of bacteriophage resistant *V. cholerae* strains. There are different ways in which *V. cholerae* can become resistant to bacteriophage infection (49, 50), one of these potential mechanisms is to mutate the O1 antigen to prevent binding of the bacteriophage to the receptor (51). Interestingly, this has been shown to result in a reduction in the ability to colonize the human intestine (52). Future studies using this model can explore whether resistant strains exhibit attenuated colonization or transmission. If eventual bacteriophage resistant mutants do arise, it would be of interest to investigate how the bacteriophage potentially evolves to circumvent this (53). If resistant strains, carrying mutations in the O1 antigen or in currently unknown resistance factors, display a decrease in the ability to colonize, this would prove as another benefit of bacteriophage therapy, where resistant strains would ultimately become less pathogenic. Thus, it could further strengthen the use of phages as a treatment for infectious diseases.

## MATERIALS AND METHODS

### Growth conditions and strains

All strains used in this study are listed in Table 1. *Vibrio cholerae* strains were cultivated under standard laboratory conditions in lysogeny broth (LB) at 30°C with shaking (25). All strains were preserved in frozen glycerol stocks and streaked on LB plates for experiments. Colonies from these plates were used for inoculation of liquid cultures. These cultures were grown overnight and used for reinoculation of liquid culture to an OD_600_ of 0.05 unless mentioned otherwise. The *Vibrio* phage ICP1 was used for all bacteriophage related experiments and was stored at 4°C in phage storage buffer (100 mM NaCl, 10 mM MgSO_4_, 10 mM Tris-HCl and 1 mM EDTA). *Vibrio cholerae* O1 biovar El Tor str. N16961 was used as the host strain for standard phage propagation procedures unless mentioned otherwise. The *V. cholerae* strains N16961-dsRED and C6706-tdtomato were provided by the Blokesch lab (Lausanne, Switzerland) and the Waldor lab (Boston, MA, USA), respectively. The ICP1 bacteriophage was provided by the Camilli lab (Boston, MA).

**Table 1:** Bacterial strains and bacteriophages used in this study.

| Strain | Genotype | Antibiotic resistance | Reference |
| --- | --- | --- | --- |
| O1 biovar El Tor str. N16961 | wt | - | (54) |
| O1 biovar El Tor str. N16961-dsRED | N16961-TndsRED | Gentamicin | (Unpublished, Blokesch lab) |
| O1 biovar El Tor str. C6706-tdtomato | C6706-tdtomato | Streptomycin | (28) |
| ICP1_2011_A; Myovirus | wt | - | (24) |

### *In vitro* phage work

#### Plaque assay

Overnight liquid culture of *V. cholerae* was diluted to an OD_600_ of 0.05 in 10 ml LB supplemented with the appropriate antibiotic (Table 1). After it had reached an OD_600_ of 0.3-0.6, 1 ml of the culture was combined with 5 mM CaCl_2_ and 50 ml soft agar (0.3% w/v agar in LB) that was cooled down to 37°C. The culture was mixed and 12.5 ml was pipetted onto prewarmed LB plates supplemented with gentamicin (10 μg/ml). The culture was solidified at room temperature for 30 minutes. A gradient of phage stock was made from 10^7^ to 10^11^ PFU/ml and 3 μl of each concentration was carefully pipetted on the plate. The plates were incubated overnight at room temperature after which the plaque counts were determined.

#### High titre phage stock preparation

A 50 ml LB culture of *V. cholerae* N16961 was grown to an OD_600_ of 0.2 followed by the addition of 10 μl ICP1 phage stock (1 × 10^9^ PFU/ml) and 5 mM CaCl_2_. After a 6 hour incubation period, the cells were centrifuged for 30 minutes at 5000 rpm (Rotations per minute) at 4°C. The supernatant was filtered through a 0.45 μm filter and incubated with phage precipitation solution (4% PEG 8000, 0.5 M NaCl) at 4°C overnight. The precipitated phages were collected by centrifugation at 3000 g for 1 hour at 4°C. Collected phages were resuspended in phage storage buffer. The phage titre was determined by plaque assay of a serial dilution as previously described.

#### *In vitro* bacteria-bacteriophage predation experiments

The *in vitro* predation experiments were performed in a similar manner to the *in vivo* infection experiments as this would allow us to closely compare the experiments. *V. cholerae* was added to the media (LB or AFW) at a concentration of 10^9^ cells per mL and diluted 1000x to mimic the amount of bacteria that were present in the wells of zebrafish larvae just before bacteriophage infection (10^6^ cell per mL). Bacteriophage ICP1 was added to the bacterial culture at roughly a MOI of 2. For the Rhine water experiment, bacteria were introduced at a higher concentration to mimic the protocol described in Silva-Valenzuela, et al. (26). Overnight grown *V. cholerae* was grown were diluted 1:1000 and grown for 8 hours at 37°C in LB. Afterwards, 20 µL was inoculated into 2.5 mL of Rhine water. Bacteriophage ICP1 was added to the culture after overnight growth in Rhine water at a MOI of 2. Serial dilution and CFU and PFU counts were performed at 0, 1, 2, 4, 7 and 24 hours.

### Cryogenic electron microscopy

Aliquots were taken at each timepoint of the growth experiment. 2 ml was concentrated to a volume of 20 µL. Quantifoil R2/2 200 mesh carbon grids (Quantifoil Micro Tools, Jena, Germany) were plasma cleaned using the Quorum Q150s glow discharger (Quorum technologies, Lewes, United Kingdoms). Aliquots of 3 μl were added to the plasma cleaned grids and subsequently plunge-frozen in liquid ethane using a Leica EMGP (Leica microsystems, Wetzlar, Germany). Blotting time was set to 1s with the chamber temperature and humidity being at 20°C and 95% respectively. Samples were transferred in grid boxes (MiTeGen, Ithaca, NY) and stored in liquid nitrogen until use. For data acquisition, grids were transferred to a Gatan 626/70 single tilt cryo-transfer holder (Gatan, Pleasanton, CA). Images were collected on Talos L120C Transmission Electron Microscope at 120kV (Thermo Fisher Scientific, Hillsboro, OR, USA).

### Zebrafish maintenance

#### Zebrafish mating and germ-free embryo rearing

Zebrafish were handled in compliance with animal welfare regulations and maintained according to standard protocols (55). The wildtype *AB/TL* line was used. All zebrafish experiments were conducted on larvae up to 5 days of age, before reaching the free-feeding stage. After the eggs were harvested, they were kept in sterile artificial fresh water (sAFW; 60 mg/L instant ocean) and treated as germ-free larvae. Germ-free larvae were generated largely as previously described, with with the following modifications (55). In brief, eggs were harvested and treated immediately with antibiotic solution (amphotericin B 250 ng/ml, kanamycin 5 μg/ml and ampicillin 100 μg/ml in sAFW), 0.1% PVP-I solution and 0.003% bleach solution resulting in a germ-free environment. The use of Gnotobiotic zebrafish medium and ZM-000 solution were replaced with sAFW. Immersion of embryos with 0.1% PVP-I solution or 0.003% bleach solution, was performed at a maximum of 1 and 5 minutes respectively. Conventionalized fish were cleaned daily and were not subjected to the protocol above. Before imaging or homogenization, larvae were anesthetized with sAFW supplemented with buffered 3-aminobenzoic acid ethyl ester to a final concentration 0.2 mg/ml (Tricaine, Sigma-Aldrich, Netherlands).

#### *In vivo* zebrafish infection

At 3 days post-fertilization (dpf), zebrafish larvae were transferred to a petri dish, with 75 larvae per well. The cells of overnight grown *V. cholerae* cultures were collected by centrifugation and resuspended in sAFW to a concentration of 10^9^ cells per ml. For the monoassociated larvae, overnight grown *Aeromonas sp*. or *Pseudomonas sp*. was centrifugated and resuspended to an OD of 0.125. 1 mL of this bacterial suspension was dissolved in a total volume of 24 mL sAFW, which corresponded to an approximate concentration of 1 × 10^6^ CFU/mL (40). The diluted bacterial suspension was introduced to the larvae 30 minutes prior to *V. cholerae* infection. After this, 1 mL of 10^9^ *V. cholerae* cells were added to the larvae in a total volume of 25 mL. The larvae were incubated with bacteria for 24 hours at 28°C. After incubation, water was removed from each well and the larvae were washed three times with sAFW to remove any excess bacteria. Imaging of the larvae would be performed 48 hours post infection at 5 dpf. The CFU of *V. cholerae* in the water of each petri dish was measured prior to the addition of bacteriophages to determine the number of phages necessary to reach the correct MOI (Table S1). An approximate estimate of the number of bacteria in the gut was based on the fluorescent signal from the imaging data. Here, we visually estimate that there are roughly 1 × 10^5^ bacteria in the intestine of a single infected larva. In total, the 75 larvae in a single petri dish sum this to roughly 7.5 × 10^6^ bacteria in the intestine of all the fish. Taking this together, the number of phages necessary to reach a MOI of 2 is 2.15 × 10^7^ PFU in each petri dish. Based on our previous *in vitro* results (Figure 1), the phage was added to the *V. cholerae* N16961-dsRED infected larvae two hours prior to imaging.

#### Fish homogenization for CFU/PFU counts

At each timepoint 5 larvae were taken from each plate and transferred to a 1.5 mL Eppendorf tube. As much water as possible was removed from the tube and 100 µL sAFW was added. The larvae were ground using a sterile pestle and a pestle motor until the suspension became homogeneous. A dilution series of this suspension was prepared and 5 µL was pipetted on both selective CFU (Gentamycin 20 µg/mL; X-gal 20 µg/mL) and PFU plates. The PFU plates were incubated at room temperature overnight and the CFU plates were incubated at 30°C overnight. This was performed at timepoints 0, 1, 2 4, 7, and 24 hours post infection with ICP1.

#### Confocal fluorescence imaging

Larvae were embedded in a drop of 1.3% (w/v) low melting point agar and carefully positioned on their lateral sides. Ethyl-m-aminobenzoate (tricaine, Sigma) was added to a final concentration of 0.2 mg/ml to tranquilize the larvae. Mounted larvae were imaged with a Leica TCS SPE confocal fluorescence microscope (Leica Microsystem, Wetzlar, Germany) using the 40x water-immersion lens. Two Z-stacks were generated for each fish at two distinct locations in the anterior intestine with the use of Leica Application Suite X (LAS X) (Figure S1).

### Serial Block Face Scanning Electron Microscopy

After fixing the material for 2 h at room temperature with 2.5% GA + 2% PFA in 0.15 M Cacodylate buffer containing 2 mM CaCl_2_, the material was washed 3 times with buffer and then placed into 2% OsO_4_ / 1.5% potassium ferrocyanide in 0.15 M Cacodylate buffer containing 2 mM CaCl_2_. The material was left for 60 minutes on ice. After washing 3 times in milliQ water, the material was placed into 1% thiocarbohydrazide for 20 m at room temperature. The material was again washed and then stained with 2% aqueous OsO_4_ for 30 m at room temperature. After washing 3 times, the material was placed into 1% Uranyl acetate for 2 h at room temperature. The material was washed with milliQ water then stained with lead aspartate for 30 m at 60°C. The material was again washed with milliQ water and then dehydrated on ice in 20%, 50% and 70% ethanol solutions for 5 m at each step. After replacing the 70% ethanol with a fresh 70% ethanol solution, the samples were kept overnight at 4°C. The next day, samples were dehydrated in 90%, 100%, 100% ethanol solutions for 5 m at each step. Next, the material was kept in dry acetone for 10 m on ice, and another 10 m in fresh dry acetone at room temperature. The material was infiltrated with 25%, 50% and 75% Durcupan ACM solution in acetone for 2 h at room temperature for each step, followed by an overnight step at room temperature in 100% Durcupan resin. The next day, the material was placed in fresh Durcupan resin for 2 h at room temperature, after which the material was flat embedded and polymerized at 60°C for 48 h.

Data were collected with a 3View2XP (Gatan Inc, Pleasanton, CA, USA) unit installed on a Zeiss Gemini 300 field emission SEM (Carl Zeiss Microscopy GmbH, Jena, Germany). The volumes were collected at 1.8 kV accelerating voltage and variable pressure at 5 Pascal. The pixel dwell time was 2 μs, with a pixel size of 10 nm and a section thickness of 75 nm.

#### Segmentation with Deep Learning and 3D visualization

The first 447 slices of the dataset were chosen to fit in an image stack with Fiji. These images were cropped around the zebrafish intestine and binned 2 times. The segmentation and the 3D visualization were done with DragonFly 2024.1. 6 slices of the dataset were manually segmented to produce the regions of interest “zebrafish_intestine”, “bacteria” and “background” (the slices 1, 80, 195, 223, 305 and 419). Then a 2D U-Net model was trained, which was pre-trained by the DragonFly team with these parameters: a data augmentation of 5 times with its parameters by default, the validation parameters by default, a patch size of 128 and batch size of 64. The training was stopped after 22 epochs, and the error of the resulting neuron network is 9%. After this training, the resulting segmentations were corrected with the 3D brush and the free-hand selection. Meshes were produced with a threshold of 50 and a sampling of 10×10×10. And the “bacteria” mesh was smoothened with 1 iteration. The workstation used has an Intel(R) Xeon(R) w7-3455 (48 CPUs), ∼2.5GHz processor with 64GB of RAM and a NVIDIA RTX A2000 graphics card with 12GB of VRAM.

#### Quantification of colonization and statistical analysis

Fiji software was used for all measurements related to the fluorescence in the intestine. Fluorescent signal of the colonization was distinguished by applying the max projection function on each Z-stack. A pixel threshold was applied between 40 and 225, the amount of pixels within these limits were counted for each max projection. Max projection was used as an indiscriminate manner of measuring colonization due to the colonizing clumps having a higher fluorescence intensity than free floating bacteria. The Z-stacks were measured in two specific areas (Figure S1) and averaged to create bar graphs for each condition and area.

Statistical analysis was done in both Prism Graphpad and excel (Microsoft Office Suite). Shapiro-Wilk test for normality and lognormality was performed to confirm lognormal distribution of fluorescence quantification. Data was log transformed to allow for parametric comparison of means. An ordinary one-way ANOVA was performed to compare the means between treatments. Error bars represent standard error of mean in log transformed data and in CFU/PFU count data. Error bars represent geometric standard error of mean in untransformed fluorescence quantification data.

## Supporting information

Movie S1: Confocal fluorescence microscopy Z-stack of an uninfected zebrafish larvae intestine

Movie S2: Confocal fluorescence microscopy Z-stack of V. cholerae N16961-dsRED infected zebrafish larvae intestine

Movie S3: Dragonfly segmentation of SBF SEM data of a V. cholerae C6706-mcherry infected zebrafish larvae intestine

Movie S4: Confocal fluorescence microscopy Z-stack of a V. cholerae N16961-dsRED and bacteriophage ICP1 infected zebrafish larvae intestine

## Data accessibility

All data are available in the main text.

## Acknowledgements

We would like to thank G. C. van der Velden, U. Nehrdich and all the others from the zebrafish facility for assisting with zebrafish larvae mating. We would also like to thank S. Munoz Sanchez for help with imaging. We are also thankful to Dr. M. Blokesch for providing the mutant *V. cholerae* strain N16961-GFP and for Dr. A. Camilli for providing the ICP1 bacteriophage.

## REFERENCES

1. M. Ali, A. R. Nelson, A. L. Lopez, D. A. Sack, Updated global burden of cholera in endemic countries. PLoS Negl Trop Dis 9, e0003832 (2015).

2. D. Hu, et al., Origins of the current seventh cholera pandemic. Proc Natl Acad Sci U S A 113, E7730–E7739 (2016).

3. L. Reveiz, et al., Chemoprophylaxis in Contacts of Patients with Cholera: Systematic Review and Meta-Analysis. PLoS One 6, e27060 (2011).

4. L. C. Dengo-Baloi, et al., Antibiotics resistance in El Tor Vibrio cholerae 01 isolated during cholera outbreaks in Mozambique from 2012 to 2015. PLoS One 12, e0181496 (2017).

5. V. Letchumanan, et al., Insights into bacteriophage application in controlling vibrio species. Front Microbiol [Preprint] (2016). [Accessed 9 September 2019].

6. J. Doss, K. Culbertson, D. Hahn, J. Camacho, N. Barekzi, A review of phage therapy against bacterial pathogens of aquatic and terrestrial organisms. Viruses [Preprint] (2017). [Accessed 9 September 2019].

7. N. K. Dutta, M. V Panse, An experimental study on the usefulness of bacteriophage in the prophylaxis and treatment of cholera. Bull World Health Organ 28, 357–60 (1963).

8. X. Wittebole, S. De Roock, S. M. Opal, A historical overview of bacteriophage therapy as an alternative to antibiotics for the treatment of bacterial pathogens. Virulence [Preprint] (2014). [Accessed 9 September 2019].

9. C. Brives, J. Pourraz, Phage therapy as a potential solution in the fight against AMR: obstacles and possible futures. Palgrave Commun 6, 1–11 (2020).

10. A. Jaiswal, H. Koley, A. Ghosh, A. Palit, B. Sarkar, Efficacy of cocktail phage therapy in treating Vibrio cholerae infection in rabbit model. Microbes Infect 15, 152–156 (2013).

11. M. Yen, L. S. Cairns, A. Camilli, A cocktail of three virulent bacteriophages prevents Vibrio cholerae infection in animal models. Nat Commun 8 (2017).

12. D. L. Runft, et al., Zebrafish as a natural host model for Vibrio cholerae colonization and transmission. Appl Environ Microbiol 80, 1710–7 (2014).

13. W. M. Spira, R. Bradley, J. L. Froehlich, “Simple Adult Rabbit Model for Vibrio cholerae and Enterotoxigenic Escherichia coli Diarrhea” (1981).

14. A. H. Meijer, H. P. Spaink, Host-Pathogen Interactions Made Transparent with the Zebrafish Model. Curr Drug Targets 12, 1000–1017 (2011).

15. H. Meijer, H. P. Spaink, Host-Pathogen Interactions Made Transparent with the Zebrafish Model. Curr Drug Targets 12, 1000 (2011).

16. Sullivan, et al., Infectious disease models in zebrafish. Methods Cell Biol 138, 101–136 (2017).

17. M. C. Gomes, S. Mostowy, The Case for Modeling Human Infection in Zebrafish. Trends Microbiol 28, 10–18 (2020).

18. Y. Senderovich, I. Izhaki, M. Halpern, Fish as Reservoirs and Vectors of Vibrio cholerae. PLoS One 5, e8607 (2010).

19. P. Noorian, M. M. Hoque, G. Espinoza-Vergara, D. McDougald, Environmental Reservoirs of Pathogenic Vibrio spp. and Their Role in Disease: The List Keeps Expanding. Adv Exp Med Biol 1404, 99–126 (2023).

20. K. C. Mitchell, P. Breen, S. Britton, M. N. Neely, J. H. Withey, Quantifying Vibrio cholerae enterotoxicity in a zebrafish infection model. Appl Environ Microbiol 83 (2017).

21. L. Plumet, D. Costechareyre, J. P. Lavigne, K. Kissa, V. Molle, Zebrafish as an effective model for evaluating phage therapy in bacterial infections: a promising strategy against human pathogens. Antimicrob Agents Chemother 68 (2024).

22. K. D. Seed, D. W. Lazinski, S. B. Calderwood, A. Camilli, A bacteriophage encodes its own CRISPR/Cas adaptive response to evade host innate immunity. Nature 494, 489–491 (2013).

23. R. Spence, et al., The distribution and habitat preferences of the zebrafish in Bangladesh. J Fish Biol 69, 1435–1448 (2006).

24. K. D. Seed, et al., Evidence of a Dominant Lineage of Vibrio cholerae-Specific Lytic Bacteriophages Shed by Cholera Patients over a 10-Year Period in Dhaka, Bangladesh. (2011). 10.1128/mBio.00334-10.

25. R. M. Martinez, C. J. Megli, R. K. Taylor, Growth and Laboratory Maintenance of Vibrio cholerae. 10.1002/9780471729259.mc06a01s17.

26. A. Silva-Valenzuela, A. Camilli, Niche adaptation limits bacteriophage predation of Vibrio cholerae in a nutrient-poor aquatic environment. 10.1073/pnas.1810138116.

27. M. Westerfield, The Zebrafish Book. A Guide for the Laboratory Use of Zebrafish (Danio rerio)., 4th Ed. (University of Oregon Press, 2000).

28. Y. A. Millet, et al., Insights into Vibrio cholerae Intestinal Colonization from Monitoring Fluorescently Labeled Bacteria. PLoS Pathog 10, e1004405 (2014).

29. Dragonfly Software for Image Processing and Data Analysis - Dragonfly. Available at: https://dragonfly.comet.tech/ [Accessed 26 August 2025].

30. J. M. Pickard, M. Y. Zeng, R. Caruso, G. Núñez, Gut Microbiota: Role in Pathogen Colonization, Immune Responses and Inflammatory Disease. Immunol Rev 279, 70 (2017).

31. A. J. Hammer, et al., Gut microbiota metabolically mediate intestinal helminth infection in zebrafish. mSystems (2024). 10.1128/MSYSTEMS.00545-24/SUPPL_FILE/MSYSTEMS.00545-24-S0004.DOCX.

32. Y. Muhammad, M. Amonov, C. Murugaiah, A. A. Baig, M. Yusoff, Intestinal colonization against Vibrio cholerae: host and microbial resistance mechanisms. AIMS Microbiol 9, 346 (2023).

33. A. Alhusayni, F. H. O. Al-Khikani, Growth of Different Bacteria on Thiosulfate Citrate Bile Salts Sucrose Agar. Journal of Marine Medical Society (2024). 10.4103/JMMS.JMMS_140_23.

34. Nabergoj, P. Modic, A. Podgornik, Effect of bacterial growth rate on bacteriophage population growth rate. Microbiologyopen 7, e00558 (2017).

35. K. G. Wiebe, B. W. M. Cook, T. J. Lightly, D. A. Court, S. S. Theriault, Investigation into scalable and efficient enterotoxigenic Escherichia coli bacteriophage production. Sci Rep 14 (2024).

36. A. Furiga, et al., Effects of Ionic Strength on Bacteriophage MS2 Behavior and Their Implications for the Assessment of Virus Retention by Ultrafiltration Membranes. Appl Environ Microbiol 77, 229 (2010).

37. S. Schink, et al., Survival dynamics of starving bacteria are determined by ion homeostasis that maintains plasmolysis. Nature Physics 2024 20:8 20, 1332–1338 (2024).

38. Y. Weng, X. R. Bina, J. E. Bina, Complete Genome Sequence of Vibrio cholerae O1 El Tor Strain C6706. Microbiol Resour Announc 10, e01301–20 (2021).

39. Lam, S. Octavia, P. Reeves, L. Wang, R. Lan, Evolution of Seventh Cholera Pandemic and Origin of 1991 Epidemic, Latin America. Emerg Infect Dis 16, 1130 (2010).

40. E. V. Koch, S. Yang, G. Lamers, J. Stougaard, H. P. Spaink, Intestinal microbiome adjusts the innate immune setpoint during colonization through negative regulation of MyD88. Nature Communications 2018 9:1 9, 1–11 (2018).

41. G. Caballero-Flores, J. M. Pickard, G. Núñez, Microbiota-mediated colonization resistance: mechanisms and regulation. Nature Reviews Microbiology 2022 21:6 21, 347– 360 (2022).

42. A. J. Silva, J. A. Benitez, Vibrio cholerae Biofilms and Cholera Pathogenesis. PLoS Negl Trop Dis 10, e0004330 (2016).

43. K. N. Wallace, S. Akhter, E. M. Smith, K. Lorent, M. Pack, Intestinal growth and differentiation in zebrafish. Mech Dev 122, 157–173 (2005).

44. M. Flores, et al., The zebrafish as a model for gastrointestinal tract-microbe interactions. (2019). 10.1111/cmi.13152.

45. A. N. Y. Ng, et al., Formation of the digestive system in zebrafish: III. Intestinal epithelium morphogenesis. Dev Biol 286, 114–135 (2005).

46. M. L. Kent, J. L. Sanders, S. Spagnoli, C. E. Al-Samarrie, K. N. Murray, Review of diseases and health management in zebrafish Danio rerio (Hamilton 1822) in research facilities. J Fish Dis 43, 637–650 (2020).

47. A. C. Borges, et al., Implementation of a Zebrafish Health Program in a Research Facility: A 4-Year Retrospective Study. Zebrafish 13, S–115 (2016).

48. J. E. Egido, A. R. Costa, C. Aparicio-Maldonado, P. J. Haas, S. J. J. Brouns, Mechanisms and clinical importance of bacteriophage resistance. FEMS Microbiol Rev 46, 1–16 (2022).

49. T. Reyes-Robles, et al., Vibrio cholerae Outer Membrane Vesicles Inhibit Bacteriophage Infection. J Bacteriol 200, e00792–17 (2018).

50. Duncan-Lowey, N. K. McNamara-Bordewick, N. Tal, R. Sorek, P. J. Kranzusch, Effector-mediated membrane disruption controls cell death in CBASS antiphage defense. Mol Cell 81, 5039-5051.e5 (2021).

51. A. Beckman, C. M. Waters, Vibrio cholerae phage ICP3 requires O1 antigen for infection. Infect Immun 91 (2023).

52. J. Nesper, et al., Characterization of Vibrio cholerae O1 El Tor galU and galE Mutants: Influence on Lipopolysaccharide Structure, Colonization, and Biofilm Formation. Infect Immun 69, 435 (2001).

53. M. Yen, A. Camilli, Mechanisms of the evolutionary arms race between Vibrio cholerae and Vibriophage clinical isolates. Int Microbiol 20, 116 (2017).

54. J. F. Heidelberg, et al., DNA sequence of both chromosomes of the cholera pathogen Vibrio cholerae. Nature 406, 477–483 (2000).

55. L. N. Pham, M. Kanther, I. Semova, J. F. Rawls, Methods for generating and colonizing gnotobiotic zebrafish. Nat Protoc 3, 1862–1875 (2008).

