## Supplemental material for "ICP1 bacteriophage treatment antagonizes colonization of the zebrafish larval intestine by *Vibrio cholerae*"

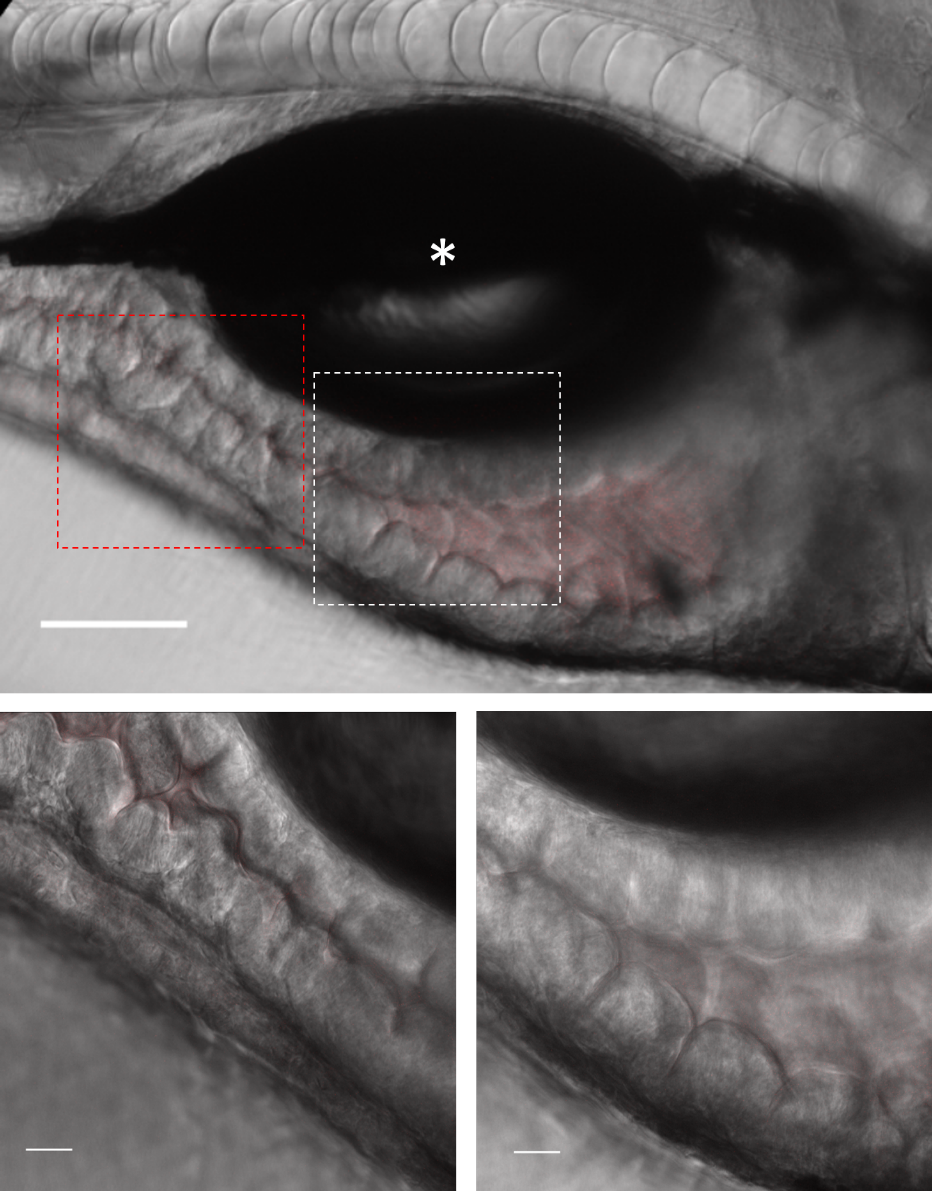

**Figure S1. Areas of interest in the zebrafish larvae intestine for the imaging of V. cholerae colonization.**

Confocal fluorescence images depicting the anterior intestine of an uninfected zebrafish larvae. The whole anterior to mid-intestine is visible (top image) with the swim bladder (asterisk) present in the middle of the image. The larva is orientated with its head to the right and its tail to left. The red background fluorescence of uninfected control larvae is visible in this image. The precise areas of interest are portrayed by the dotted boxes. Single slice images of Z-stacks of both the mid-intestine (red box) and the anterior intestine (white box) can be seen below. Scale bars represent 100 μm (top image) and 20 μm (bottom images) respectively.

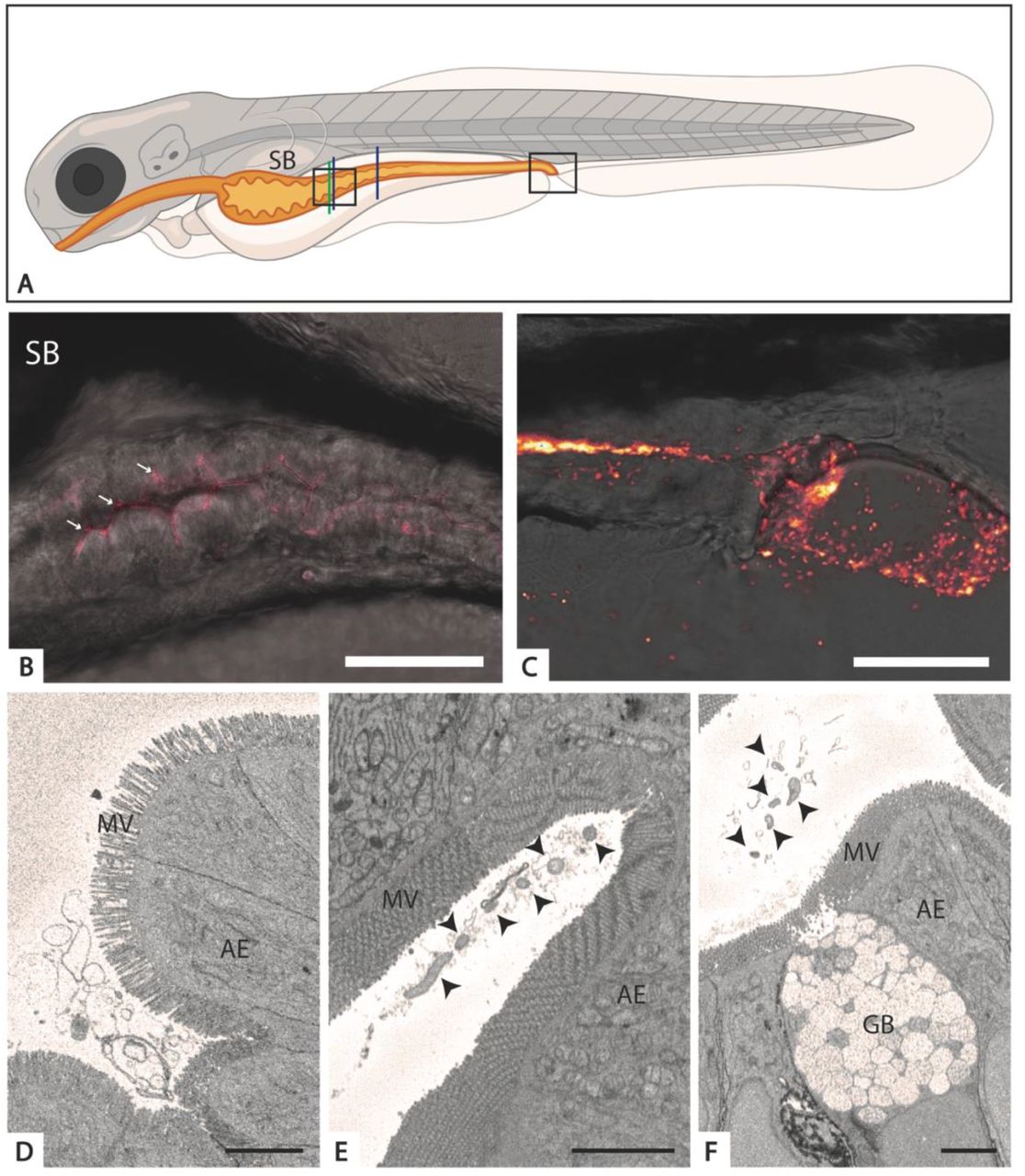

**Figure S2. Colonization of the mid-intestine of the zebrafish larvae by *V. cholerae* C6706-tdTomato**

Fluorescent imaging of the mid-intestine. The white arrows indicate areas of colonization. Scale bar represents 75 μm.

**C6706-tdTomato**

**Figure S3. Infection and clearance of *V. cholerae* N16961-DSRED by ICP1 in germ-free zebrafish larvae 4 hours post bacteriophage infection.**

Single slice images of confocal fluorescence Z-stacks showing the anterior intestine of germ-free zebrafish larvae both untreated (top) and treated by ICP1 (bottom). The intestine was imaged directly below the swim bladder (left) and at the mid-intestine (right). Scale bars represent 20 μm.

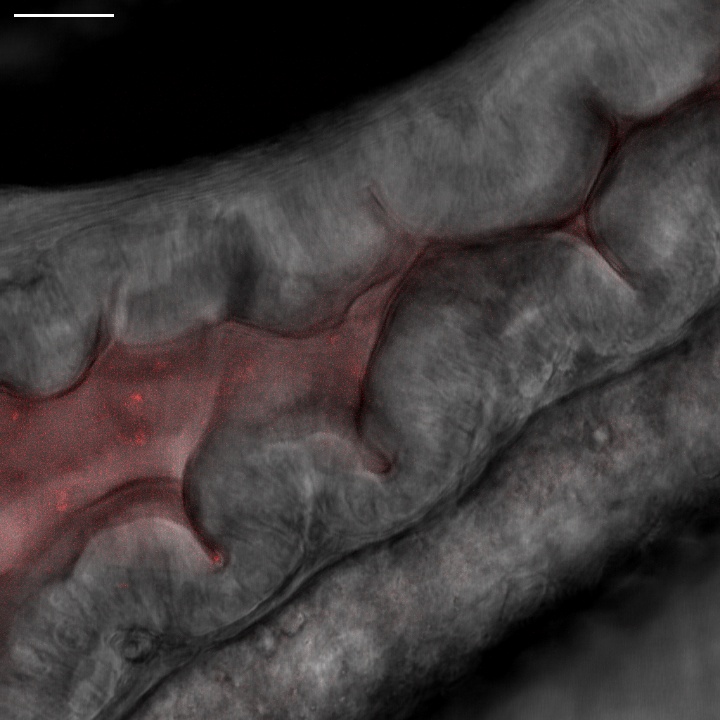

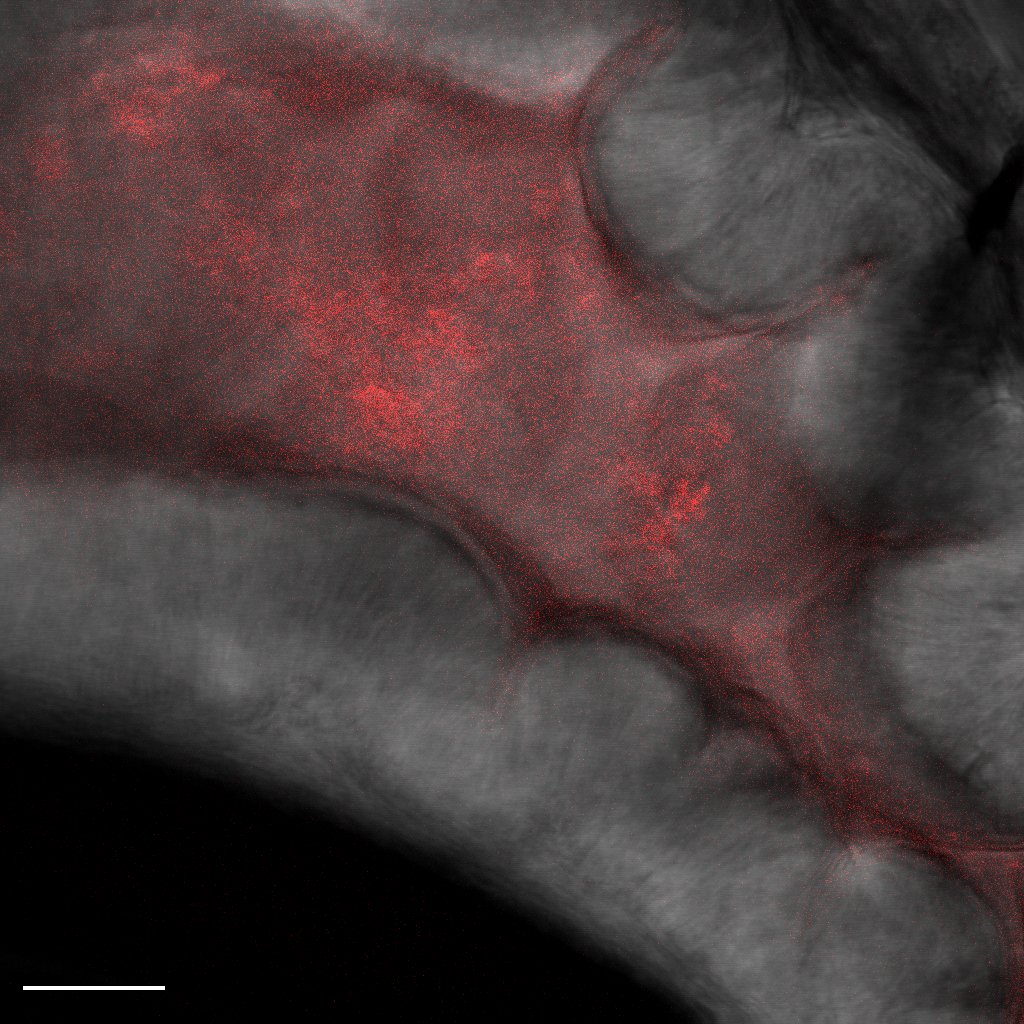

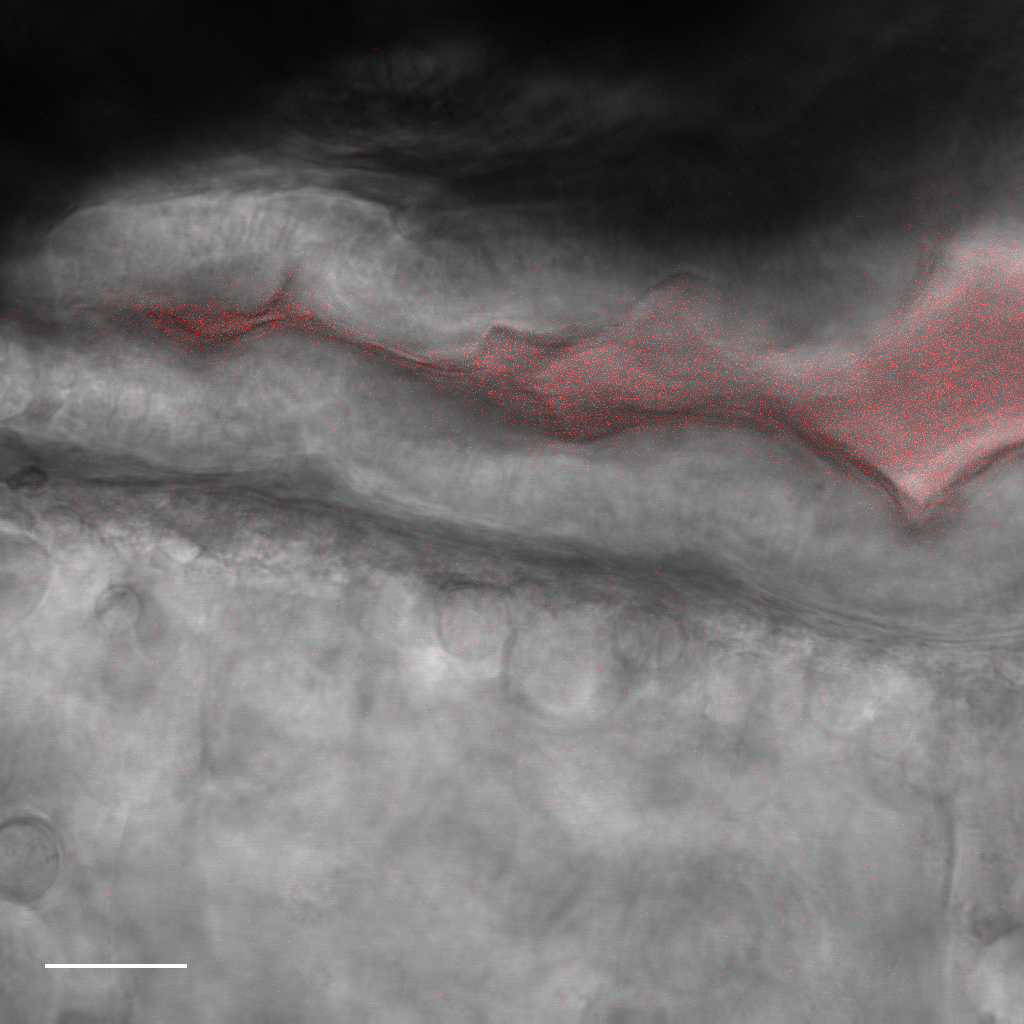

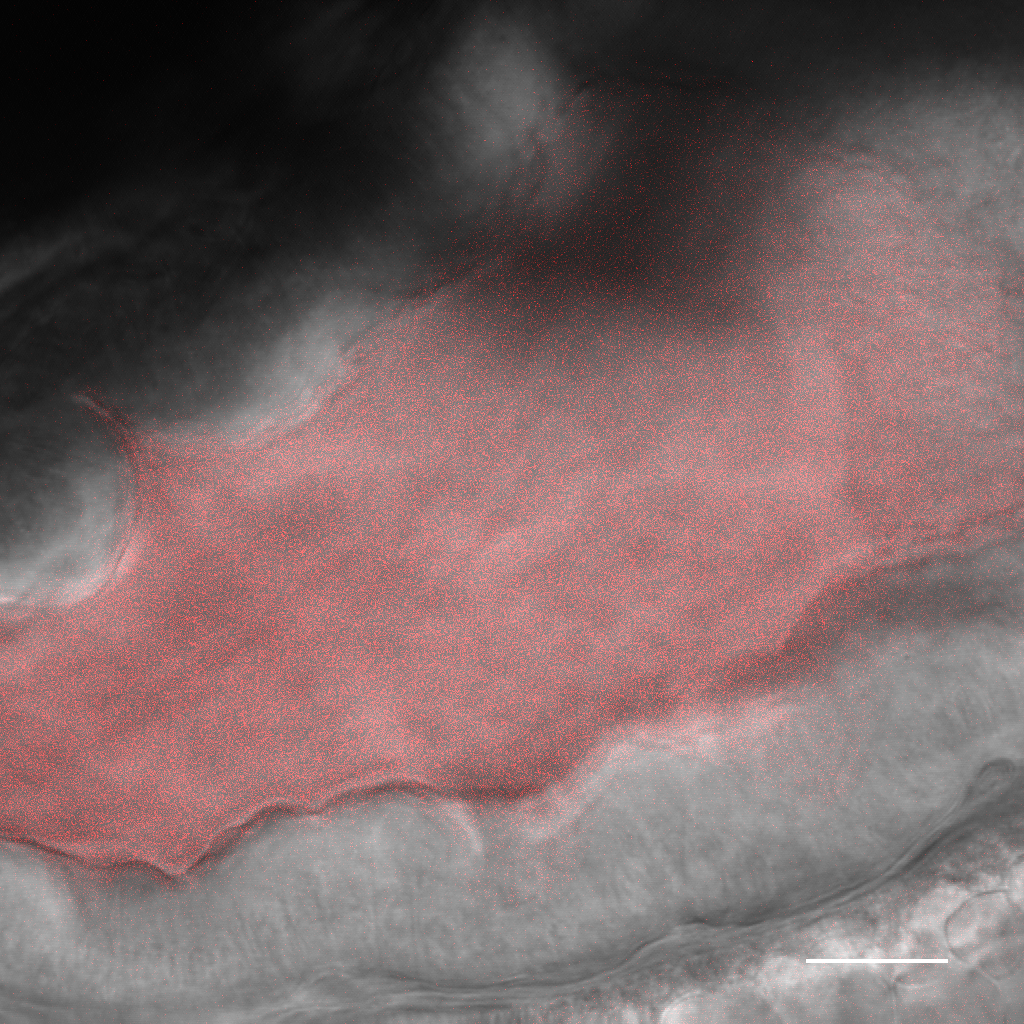

**N16961-DSRED**

**N16961-DSRED**

**N16961-DSRED**

**N16961-DSRED**

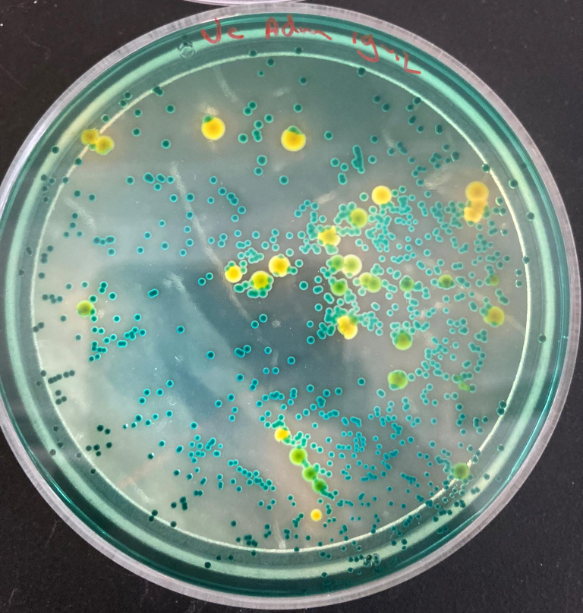

**Figure S4. TCBS plate displaying the two colony morphologies of *Aeromonas sp.* (yellow) and *Pseudomonas sp.* (green)**

**Figure S5. Fluorescent quantification of *V. cholerae* N16961 in zebrafish larvae intestine**

(A) Representative images depicting the fluorescent pixel quantification workflow. A single slice image fluorescent image is depicted (left). The max intensity projection of this image is shown (middle) and a lower threshold of 40 is applied (right). All the pixels above this threshold are quantified. (B) Graphs depicting the geometric mean of the quantified pixels of colonizing bacteria in the anterior (left) and mid-intestine (right) for both germ-free (top) and conventionalized zebrafish larvae (bottom). Error bars represent geometric standard deviation.

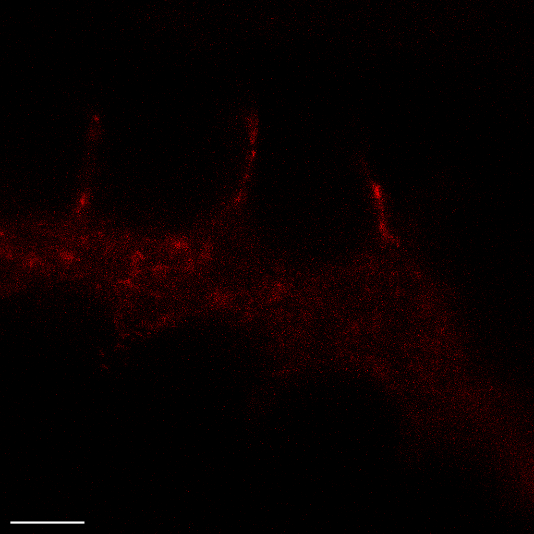

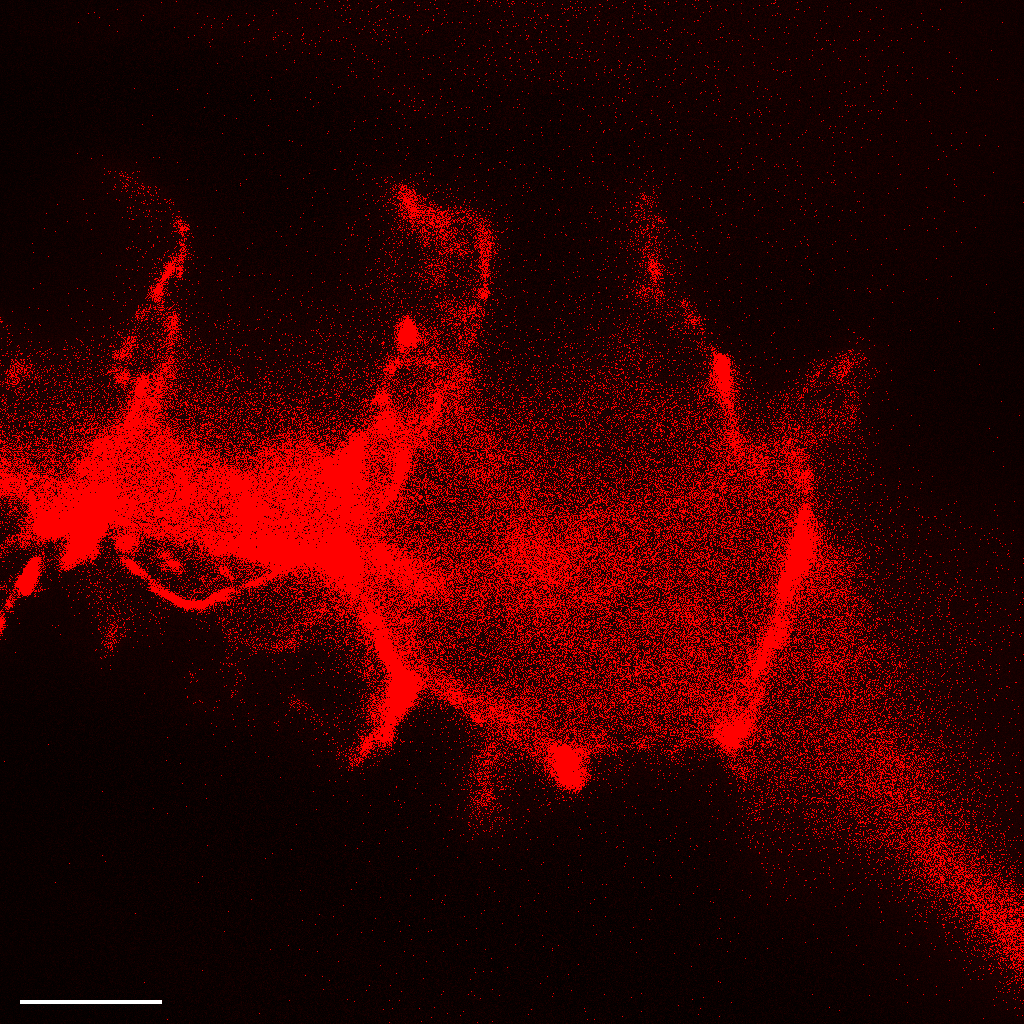

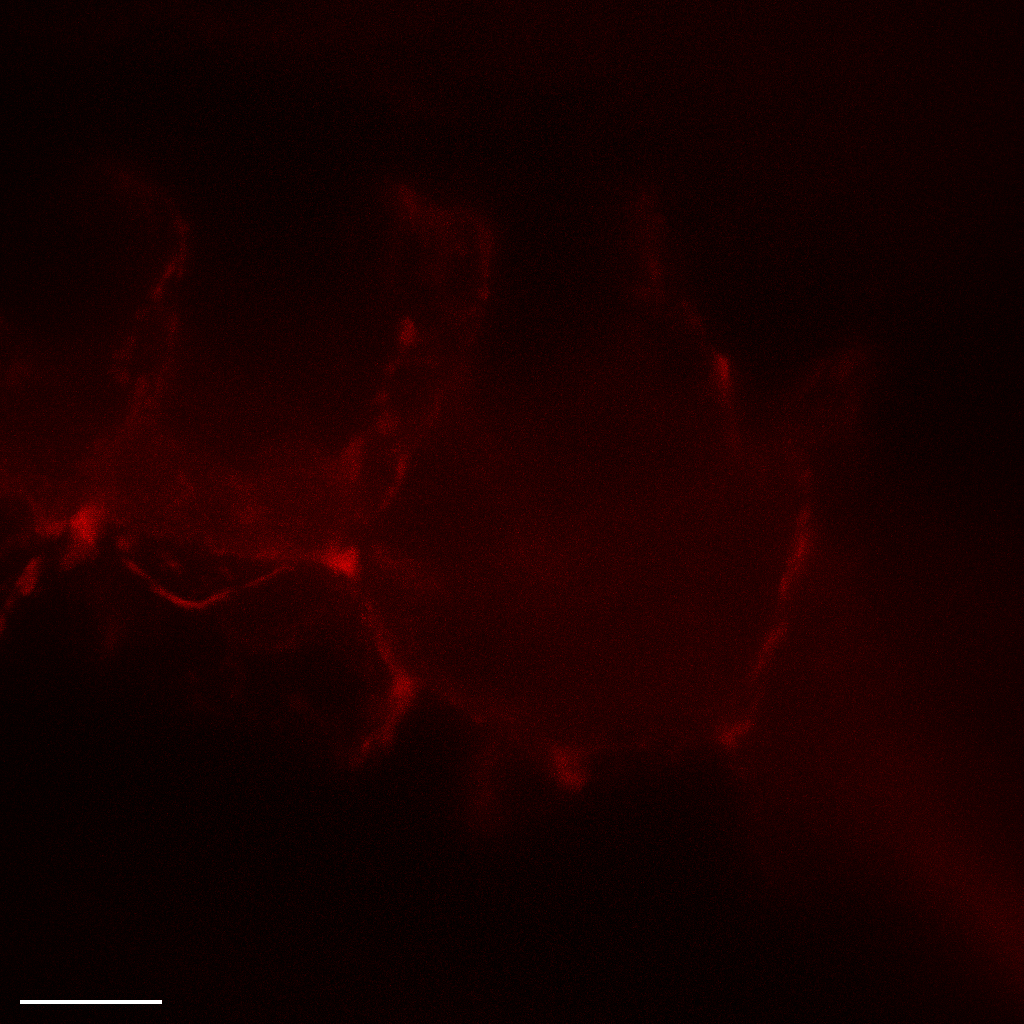

**N16961-DSRED**

**N16961-DSRED**

**N16961-DSRED**

**A.**

**B.**

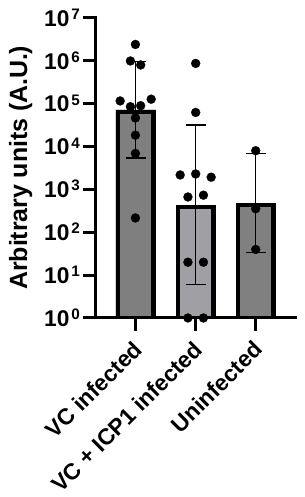

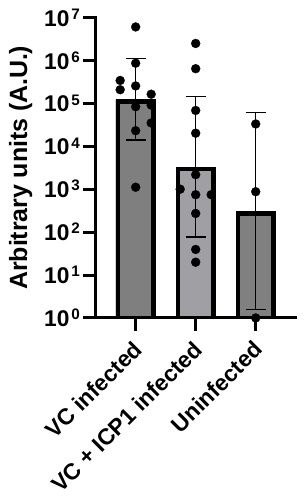

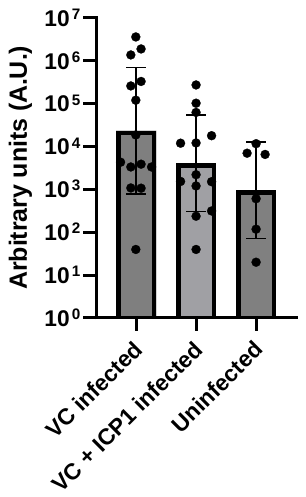

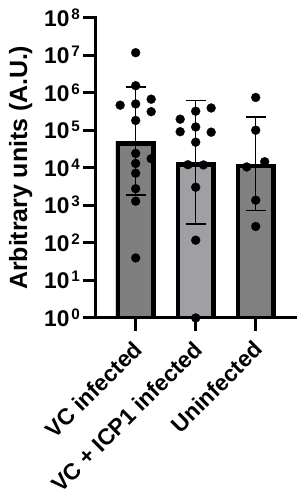
